# Exploiting synergistic interactions of *Medicago sativa* L. and *Paraburkholderia tropica* for enhanced biodegradation of diesel fuel hydrocarbons

**DOI:** 10.1101/2021.03.30.437699

**Authors:** Michael O. Eze, Volker Thiel, Grant C. Hose, Simon C. George, Rolf Daniel

**Author notes:** These authors contributed equally to this work (Simon C. George, Rolf Daniel).

## Abstract

The biotechnological application of microorganisms for rhizoremediation of contaminated sites requires the development of plant-microbe symbionts capable of plant growth promotion and hydrocarbon degradation. Studies focusing on microbial consortia are often difficult to reproduce, thereby necessitating the need for culturable single bacterial species for biotechnological applications. Through genomic analyses and plant growth experiments, we examined the synergistic interactions of *Medicago sativa* L. and *Paraburkholderia tropica* for enhanced remediation of diesel fuel-contaminated soils. Comparative genomics revealed strong potential of *P. tropica* for plant growth-promotion, chemotaxis and motility, root nodulation and colonization, and diesel fuel degradation. Plant growth experiments confirmed that *P. tropica* thrived in the contaminated soils and effectively enhanced *M. sativa* growth. Geochemical analysis showed that the *M. sativa* + *P. tropica* treatment resulted in an efficient degradation of diesel fuel hydrocarbons within two months, offering great prospects for enhanced biodegradation of organic pollutants.

## INTRODUCTION

The quest for energy to meet the growing global demand has depended largely on petroleum and other fossil fuels. Consequently, petroleum spillage, either through human error or equipment failure has plagued the environment for decades^1^. For example, in 2010, the Deepwater Horizon oil spill discharged more than 700 million litres of South Louisiana crude oil into the Gulf of Mexico, resulting in the largest marine oil spill in U.S. history and the second largest in the world^2, 3^. While large-scale marine spills often make headlines, the majority of petroleum spills occur on land, with significant human health and ecological impacts^4^

Remediation of petroleum contaminated sites is essential to mitigating the human and ecological risks. Remediation strategies are classified as either *ex situ* or *in situ*^5^. *Ex situ* techniques involve the excavation and relocation of contaminants for off-site treatments and, as a result, are expensive and environmentally unfriendly. In contrast, *in situ* remediation involves the on-site treatment of contaminants, which is typically more eco-friendly and 50 to 80% cheaper than traditional methods such as excavation and landfill incineration^6^.

Phytoremediation, that is, the use of plants to remediate contaminated sites, is a cost-effective method for *in situ* remediation. This technique relies on the use of plant interactions (physical, biochemical, biological, chemical, and microbiological) to remediate the toxic effects of pollutants^7, 8^. This technique is based on carefully selected plants with fibrous roots that serve as a natural host for hydrocarbon-degrading microorganisms. The extensive rooting systems allow air to enter the rhizosphere, thus serving as a natural bioventing system, leading to increased biodegradation of pollutants. Notwithstanding the merits of this technique, the slow metabolic activity of the hydrocarbon-degrading microorganisms leads to longer remediation times, limiting the application of the technique. To address this shortfall, there is growing interest in the isolation of microbial consortia usable as inocula for contaminant degradation.

In addition to their hydrocarbon-degrading capacities, some of the inoculated microbes can potentially enhance the growth of host plants through processes such as nitrogen fixation, and phosphate and potassium solubilization^9^ In turn, the root exudates released by the host plants provide nutrition to the associated bacteria, enabling continuous biodegradation of contaminants. This synergistic relationship has been described as the ecological driver of rhizoremediation^10^. Therefore, when carefully applied, bioaugmentation offers great potential for effective reclamation of hydrocarbon contaminated sites.

While a number of studies have been carried out on the use of microbial consortia for rhizoremediation^11, 12^, few studies have been performed with single bacterial isolates. This is a major challenge to the adoption of microbially-enhanced rhizoremediation since the replication of consortia is always difficult. Hence, there is an urgent need to isolate single culturable bacterial species for biotechnological applications. Therefore, the aims of this study were to isolate potential plant growth-promoting and hydrocarbon-degrading single bacterial species, and examine their capacity to enhance the rhizodegradation of diesel fuel hydrocarbons. Specifically, this study is the first attempt to examine the synergistic interactions of *Medicago sativa* L. and *Paraburkholderia tropica* for enhanced rhizoremediation of diesel fuel-contaminated soils. This is highly relevant considering recent studies have shown that *Paraburkholderia* strains can potentially promote plant growth and/or degrade contaminants^13, 14^ Additionally, our earlier studies revealed that *M. sativa* has high tolerance to petroleum hydrocarbons, relative to other plant species^15^, thereby making it an ideal plant to investigate plant-microbe synergy for biodegradation. By combining genome studies of *P. tropica* with a pot-based rhizodegradation experiment, we demonstrate that synergistic interactions between *M. sativa* and *P. tropica* promotes rhizodegradation of petroleum hydrocarbons.

## RESULTS

### Genome Sequencing

The sequencing of the three genomes resulted in a total of 4,228,774 gene sequences. After processing, a total of 3,450,275 paired-end reads and 356,023 unpaired reads were retained and assembled. Assembly resulted in 575 scaffolds (Supplementary Table S1). The average sequencing depth was approximately 30x. The three genomes were taxonomically classified as *Acidocella facilis, Burkholderia* sp. and *Paraburkholderia tropica*.

### Functional Analysis of the Genomes

#### Plant growth-promotion, chemotaxis, motility and root colonization

Functional analysis of the three genomes revealed that *P. tropica* has the greatest potential for plant growth promotion, with 36, 31, 4, 2 and 2 coding DNA sequences (CDSs) putatively involved in nitrogen fixation, phosphate solubilization, pyrroloquinoline quinone synthesis, siderophore transport and indoleacetic acid synthesis respectively (Supplementary Table S2). Key genes involved in these processes include the *nifAUQ, fixABJL, acpP, otsB, gph, plc, pqqBCDE, entS* and *iaaH* genes (Table 1). In comparison, the genomes of the other bacterial isolates had far fewer (24-44) CDSs putatively involved in plant growth promotion, with *nifAQ, phoD, entS* and *iaaH* genes missing in the genome of *A. facilis* (Supplementary Table S2).

**Table 1.** Key genes in *Paraburkholderia tropica* genome that are potentially involved in plant growth promotion, bacterial chemotaxis, motility, root nodulation and colonization (See Supplementary Table S2 for more details).

| Genes | CDSs | Enzyme | Function |
| --- | --- | --- | --- |
| ***Plant growth promotion*** |  |  |  |
| *nifA* | 2 | Nif-specific regulatory protein | Nitrogen fixation |
| *nifUQ* | 2 | Nitrogen fixation proteins NifU, NifQ | Nitrogen fixation |
| *fixAB* | 10 | Electron transfer flavoprotein | Nitrogen fixation |
| *fixJL* | 22 | Two-component system, LuxR family | Nitrogen fixation |
| *acpP* | 4 | Acyl carrier protein | Phosphate solubilization |
| *otsB* | 3 | Trehalose 6-phosphate phosphatase | Phosphate solubilization |
| *gph* | 6 | Phosphoglycolate phosphatase | Phosphate solubilization |
| *plc* | 9 | Phospholipase C | Phosphate solubilization |
| *pqqBDE* | 3 | Pyrroloquinoline quinone biosynthesis proteins B, D, E | PQQ synthesis |
| *pqqC* | 1 | Pyrroloquinoline-quinone synthase | PQQ synthesis |
| *entS* | 2 | MFS transporter, ENTs family, enterobactin exporter | Siderophore transport |
| *iaaH* | 2 | Indoleacetamide hydrolase | IAA synthesis |
| ***Bacterial chemotaxis, motility, and root colonization*** |  |  |  |
| *cheABCDVWXYZ* | 20 | Two-component chemotaxis protein | Bacterial chemotaxis |
| *wspBDEF* | 4 | Two-component chemotaxis protein | Bacterial chemotaxis |
| *mcp*, *tsr*, *tar*, *trg*, *tap*, *wspA* | 47 | Methyl-accepting chemotaxis protein | Bacterial chemotaxis |
| *flg*, *flh*, *fli* | 52 | Flagellar proteins | Bacterial motility |
| *nodD* | 2 | Nod-box dependent transcriptional activator | Root nodulation |
| *tadB* | 2 | Tight adherence protein B | Plant colonization |
| *tadC* | 2 | Tight adherence protein C | Plant colonization |

PQQ: Pyrroloquinoline quinone; IAA: Indole-3-acetic acid.

#### Hydrocarbon-degrading potential

Functional analysis revealed that *P. tropica* contains genes putatively involved in hydrocarbon degradation (Table 2). While the three isolates are potentially able to degrade petroleum hydrocarbons, there were some specific differences in their metabolic capabilities. For example, the genomes of *P. tropica* and *Burkholderia* sp. encode for greater numbers of enzymes putatively involved in the degradation of *n*-alkanes (especially long-chain *n*-alkanes) and cycloalkanes than the *Acidocella* genome (Supplementary Table S3). This is crucial for diesel fuel degradation since the chemical composition of diesel fuel is 75% saturated hydrocarbons (predominantly long-chain *n*-alkanes and cycloalkanes)^16, 17^

**Table 2.**
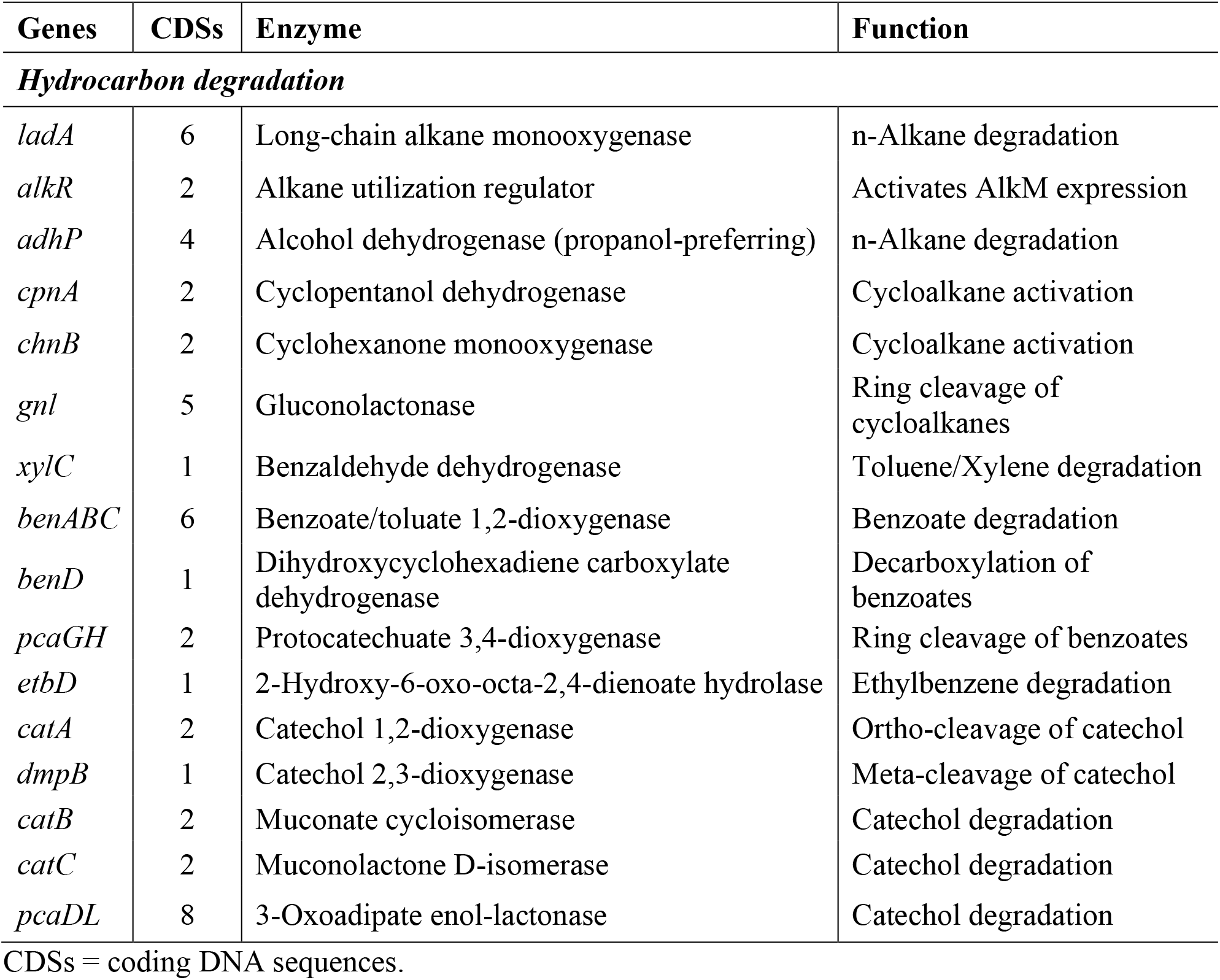
Key genes in the *Paraburkholderia tropica* genome that putatively encode for hydrocarbon-degrading enzymes.

The degradation of aromatic hydrocarbons proceeds through initial activation by monooxygenases, followed by ring cleavage of catechols. While genes putatively encoding for monooxygenases and dehydrogenases were more abundant in the genome of *P. tropica*, genes encoding for ring-activating dioxygenases (such as benzoate/toluate 1,2-dioxygenase) were more abundant in the *A. facilis* genome (Supplementary Table S3). In addition, putative DNA sequences encoding for ortho-cleavage of catechol (*catABC* and *pcaDL* genes) were more abundant in the *P. tropica* and *Burkholderia* sp. genomes than in the genomes of *A. facilis* (Supplementary Table S3).

### Effect of Bacterial Inoculation on Plant Growth and Biomass Production

The inoculation of *M. sativa* with *P. tropica* resulted in increased growth rate and biomass production (Figure 1). The analysis of variance (ANOVA) revealed that the mean dry biomass (± standard error, SE) produced by plants inoculated with *P. tropica* (6.74 ± 0.06 g) was significantly higher than, and significantly different (*p*< 0.001) from that of the uninoculated plants (3.38 ± 0.07 g) (Figure 1b). Statistical analysis of growth (in terms of mean shoot height per time) revealed that the inoculated *M. sativa* plants exhibited greater relative growth rate (0.109 ± 0.002 cm/day at the point of inflection) than the uninoculated plants (0.092 ± 0.004 cm/day) (Figure 1c).

**Figure 1.**
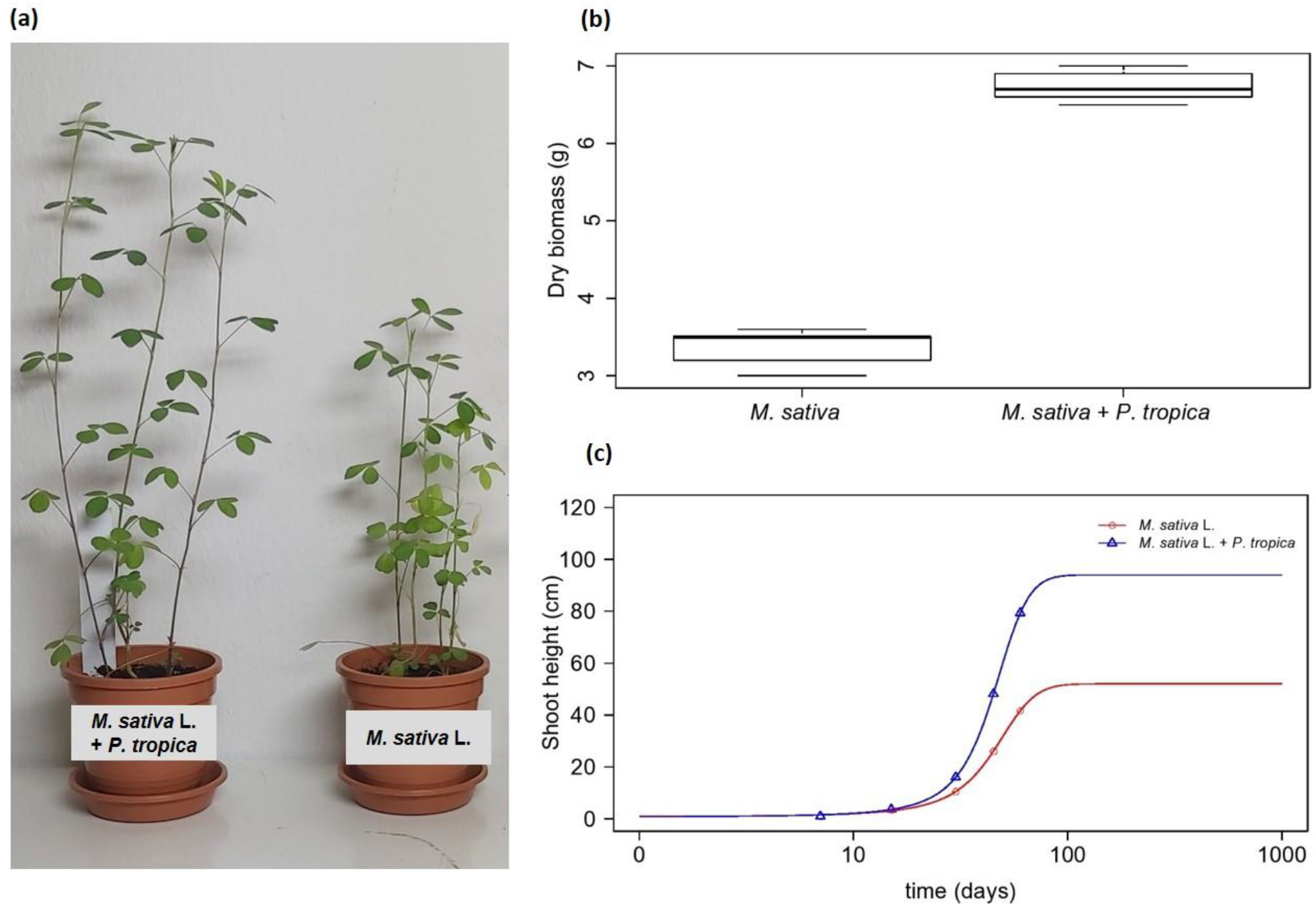
(a) Enhancement of growth and biomass production of *M. sativa* plant by *P. tropica* inoculation. (b) Boxplot showing significant differences in dry biomass of inoculated and uninoculated plants. (c) 3-parameter logistic model showing differential growth rates (cm/day) of *M. sativa* inoculated with *P. tropica* vs. uninoculated control.

### Organic Geochemical Analysis of Biodegradation

#### Total petroleum hydrocarbons

Geochemical analysis of residual hydrocarbons revealed that the inoculation of contaminated soils with the *P. tropica* isolate significantly enhanced hydrocarbon degradation and reclamation of the diesel fuel-contaminated soils (Figure 2). As shown in the gas chromatograph-mass spectrometry (GC-MS) chromatograms, nearly all the distinctive diesel fuel hydrocarbons, including C_10_–C_25_ *n*-alkanes, branched alkanes, cyclic alkanes and aromatic hydrocarbons (approximately 96%) were degraded in the planted and inoculated soils (Figure 2a). Similar results were obtained with the replicates (Supplementary Figure S1). The mean total petroleum hydrocarbons (± SE) at the beginning of the experiment (T0) was 4.13 ± 0.04 g/kg. At the end of the experimental period, the greatest decrease in residual total petroleum hydrocarbons was observed in the “planted and inoculated soils” (Soil+*M.sativa+P.tropica*), while the least decrease occurred in the “unplanted and uninoculated soils” (Soil at T60) (Figure 2b). This is an indication of greatest biodegradation in the “Soil+*M.sativa*+*P.tropica*” treatment, a claim further supported by the huge UCM observed under this treatment (Figure 2c). Preparation blanks ruled out possible cross contamination of hydrocarbons among the samples (Supplementary Figure S2).

**Figure 2.**
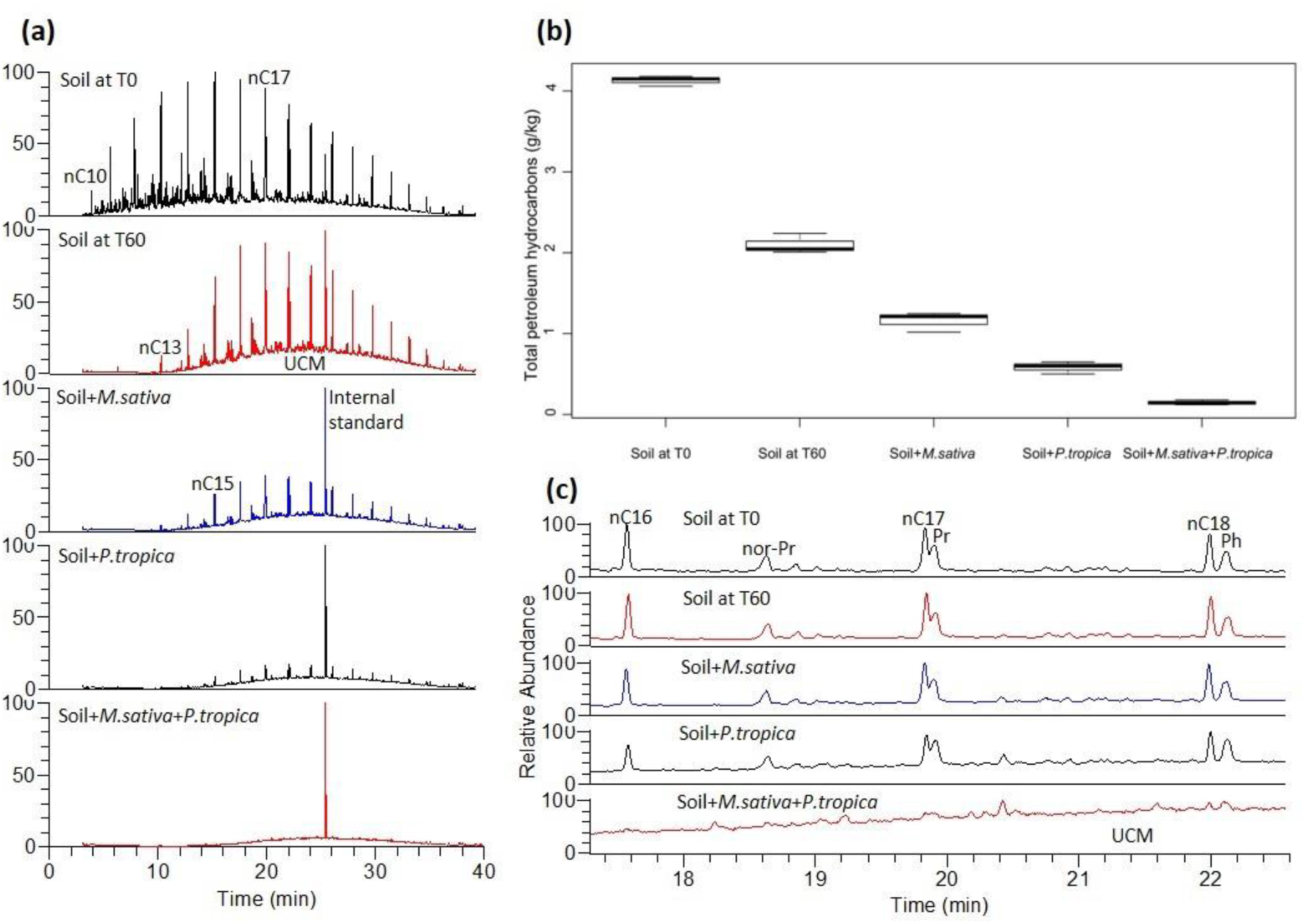
(a) Partial total ion chromatograms showing the initial total hydrocarbons in the contaminated soils and the residual hydrocarbons after 60 days under different treatments. The chromatograms can be directly compared with reference to the internal standard peak (the same amount was added to all samples). (b) Boxplot showing the mean values of residual total petroleum hydrocarbons for the different treatments. (c) Partial total ion chromatograms showing preferential biodegradation of nC16, nC17 and nC18 relative to nor-pristane (nor-Pr), pristane (Pr), and phytane (Ph) with increasing biodegradation among the different treatments. “Soil at T0”: contaminated soils at the beginning of experiment; “Soil at T60”: unplanted and uninoculated soils after 60 days; “Soil+*M.sativa*”: soils planted with *M. sativa* L.; “Soil+*P.tropica*”: soils inoculated with *P. tropica*; “Soil+*M.sativa+P.tropica*”: soils planted with *M. sativa* L. and inoculated with *P. tropica;* UCM: unresolved complex mixture.

#### Biodegradation parameters

The highest values of nC17/Pr, nC18/Ph, nC16/nor-Pr and total petroleum hydrocarbons/unresolved complex mixture (TPH/UCM) (1.34, 1.39, 2.65 and 1.82 respectively) were observed in the aged initial contaminated soil (Soil at T0) (Table 3), followed by the unplanted and uninoculated contaminated soil at the end of the experimental period (Soil at T60). The lowest values of these ratios, indicating most intense biodegradation, were found in the planted and inoculated soils (Soil+*M.sativa+P.tropica*). The nC17/Pr, nC18/Ph, nC16/nor-Pr and TPH/UCM ratios for the treatment “Soil+*P.tropica*” were smaller than that of “Soil+*M.sativa*” treatment (Table 3). This is an indication that the inoculation of diesel fuel-contaminated soils with *P. tropica* alone resulted in greater hydrocarbon degradation than simple phytoremediation using *M. sativa*. This is also in agreement with the results of total residual hydrocarbon measurements (Figure 2b).

**Table 3.**
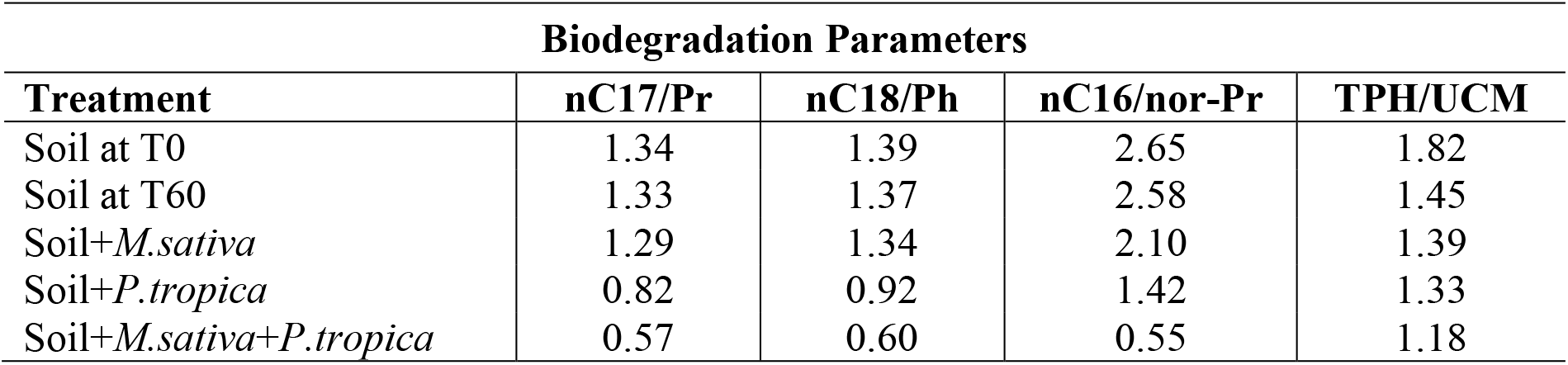
Biodegradation parameters (ratios) for the different treatments.

Tukey’s pairwise comparisons showed that the mean total petroleum hydrocarbons (g/kg) in the soils at the end of the experiment (Soil at T60) were significantly different from the initial mean concentration (Soil at T0), indicating that biodegradation occurred in the treatments (Supplementary Table S4). It also revealed that the residual hydrocarbon concentrations differed significantly between each treatment (Figure 2b; Supplementary Table S4). The greatest percentage of biodegradation (96%) observed in the planted and inoculated samples (Soil+*M.sativa+P.tropica*) were the result of the synergistic actions of *M. sativa* and *P. tropica*.

#### Biodegradation of polycyclic aromatic hydrocarbons

Molecular analysis of residual polycyclic aromatic hydrocarbons such as alkylnaphthalenes and alkylphenanthrenes revealed that the combined application of *M. sativa* and *P. tropica* resulted in complete biodegradation of these otherwise recalcitrant hydrocarbons (Figure 3). When compared to other treatments, “*M. sativa*+*P. tropica*” treatment appeared to be the most-effective approach for biodegradation of these organic pollutants (Supplementary Figure S3).

**Figure 3.**
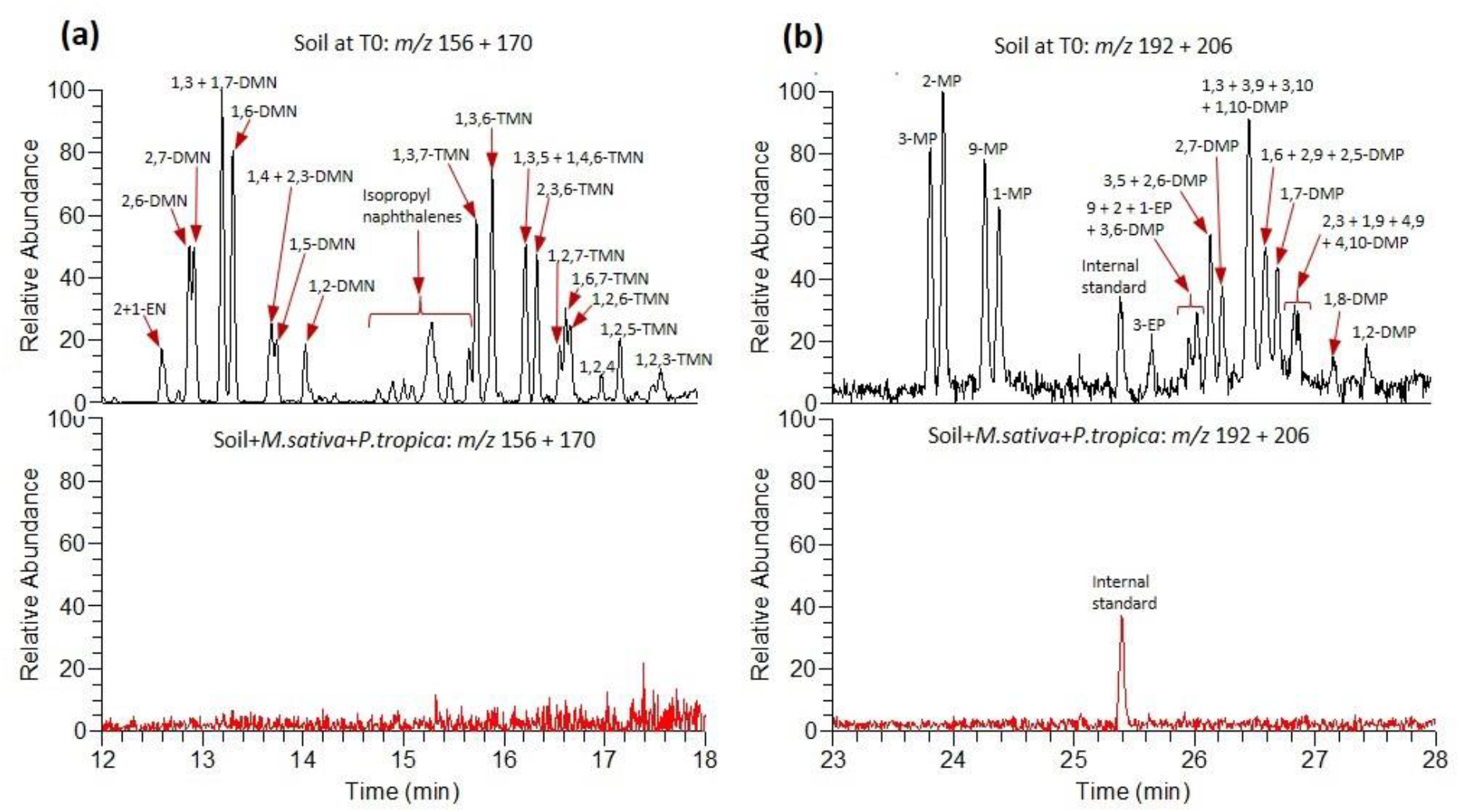
Partial (a) *m/z* 156 + 170 and (b) *m/z* 192 + 206 mass chromatograms of contaminated “soil at T0” and “planted and inoculated soil” at the end of the experiment, showing effective biodegradation of alkylnaphthalenes and alkylphenanthrenes by the combined actions of *M. sativa* and *P. tropica*. EN: ethylnaphthalene; DMN: dimethylnaphthalene; MP: methylphenanthrene; EP: ethylphenanthrene; DMP: dimethylphenanthrene. Numbers denote positions of alkylation.

#### 16S rRNA Analysis of the Residual Soils

The relative abundances of *Paraburkholderia* in the uninoculated soils (both planted and unplanted) were less than 1%. In contrast, the relative abundance of *Paraburkholderia* in the inoculated soil was approximately 5%, making it the fourth most-abundant bacterial genera in the inoculated soils (Supplementary Figure S4). This indicates that the inoculated microbes prospered in the rhizosphere.

## DISCUSSION

The results of whole genome analysis revealed that *P. tropica* is a potential plant growth-promoting bacterium (Table 1). Functional analysis of the genomes of the three isolates revealed that *P. tropica* has the highest potential for plant growth promotion. For example, while *A. facilis has* 11 coding DNA sequences (CDSs) potentially involved in nitrogen uptake processes, *P. tropica* has 36 CDSs potentially involved in nitrogen metabolism (Supplementary Table S2). Similarly, *P. tropica* isolates possess the highest number of genes encoding phosphatases, siderophore exporter and indoleacetic acid synthase. The potential of *P. tropica* to enhance plant growth through these processes have also been documented in few recent studies^14, 18^.

The genome of *P. tropica* revealed other important genes involved in chemotaxis, motility, and root colonization. These processes are highly important if the inoculated isolate is to prosper in the rhizosphere. Chemotaxis proteins identified in the genome include 24 genes encoding two-component systems (*cheABCDVWXYZ* and *wspBDEF* genes) and 47 methyl-accepting chemotaxis proteins. This is about 2–4 times the number present in the genomes of the other bacteria (Supplementary Table S2). We also identified 52 flagellar biosynthesis proteins belonging to the *flg, flh* and *fli* genes. These genes are crucial for bacterial motility. For example, Böhm, Hurek and Reinhold-Hurek^19^ found that the deletion of genes involved in motility in the endophyte *Azoarcus* sp. prevented their twitching, motility and colonization of rice plants. The *nod* and *tad* genes are vital for root nodulation and plant colonization. The *nodD* gene has previously been associated with root nodulation in *Rhizobium*^20^. Similarly, previous in silico and experimental studies of other plant growth promoting bacteria have linked the tight adherence *(tad)* systems to plant attachment and colonization^21–23^. The presence of more genes encoding for plant growth-promoting processes, and for chemotaxis, motility, and root colonization in *P. tropica* than in the other bacteria, strongly suggests a possible advantage in rhizoremediation.

Diesel fuel is phytotoxic to most plant species, and therefore often has negative effects on plant growth and biomass production^15^. The inoculation of *M. sativa* with *P. tropica* in this study resulted in increased growth rate and biomass production of *M. sativa*. The mean dry biomass of inoculated plants was more than twice that of the uninoculated plants (Figure 1). Additionally, during the 60-day experimental period, the inoculated plants attained an average height of 80 cm, in contrast to approximately 42 cm for the uninoculated plants. These results are not surprising in view of the strong plant growth-promoting potential of *P. tropica* as shown by the genome analysis.

Rhizoremediation of petroleum-contaminated soils also depends on the ability of the inoculated bacteria to degrade hydrocarbons. The examination of the three genomes revealed that all of them are potentially able to utilize aliphatic and aromatic hydrocarbons as carbon and energy sources. Key hydrocarbon-degrading enzymes present in the genome of *P. tropica* include long-chain alkane monooxygenase, cyclohexanone monooxygenase, benzoate/toluate 1,2-dioxygenase, protocatechuate 3,4-dioxygenase, catechol 1,2-dioxygenase and catechol 2,3-dioxygenase (Table 2). Among the three genomes, *P. tropica* genome possesses more genes potentially involved in long-chain alkane (*ladA* and *alkR* genes) and cycloalkane (*cpnA, chnB* and *gnl* genes) degradation. Since diesel fuels and similar low-volatile petroleum distillates are composed predominantly of aliphatic hydrocarbons (approximately 75%)^16, 17^, the presence of more high-molecular weight aliphatic hydrocarbon-degrading genes in *P. tropica* than in other species suggests stronger potential for the biodegradation of diesel spills. The presence of ring-cleavage dioxygenase such as catechol 1,2-dioxygenase and catechol 2,3-dioxygenase in the genome of *P. tropica* also indicates its potential to metabolise intermediate products of aromatic hydrocarbon degradation. In a recent study, a particular strain of *Paraburkholderia aromaticivorans* isolated from a petroleum-contaminated soil was found capable of degrading naphthalene and BTEX (benzene, toluene, ethylbenzene and xylene)^24^ In addition, *Paraburkholderia* isolate exhibited higher growth rate in neutral pH (as determined by OD_600_ values) than either the *Acidocella* or the *Burkholderia* isolate, indicating potential wider application of *P. tropica* for rhizoremediation than the other species. Conversely, the adaptability of *Acidocella* to acidic conditions can be exploited for the remediation of sites contaminated with metals and organic pollutants such as mining sites^25–27^.

Geochemical analysis of biodegradation revealed that *M. sativa*+*P. tropica* treatment significantly enhanced the biodegradation of diesel fuel hydrocarbons, resulting in 96% rhizodegradation of the total petroleum hydrocarbons (Figure 2). Natural attenuation led to only 49% degradation. In comparison, *M. sativa* alone, *P. tropica* alone, and *M. sativa*+*P. tropica* treatments resulted in 72%, 86% and 96% biodegradation respectively. Biodegradation parameters revealed that the removal of petroleum hydrocarbons in the different treatments came from biodegradation. These parameters (nC17/Pr, nC18/Ph, nC16/nor-Pr and TPH/UCM) followed the expected trends, with *n*-alkanes more biodegradable than branched alkanes, and the UCM more recalcitrant than TPH. The parameters indicate that the highest degree of biodegradation among the different treatments occurred in “planted and inoculated” soils (Table 3). *Tukey*’s pairwise comparisons confirmed these results and also indicated that the results of biodegradation were significantly different among the different treatments.

Although the chemical composition of diesel fuel is predominantly *n*-alkanes, branched alkanes and cycloalkanes, it also contains approximately 25% aromatic hydrocarbons such as alkylbenzenes, naphthalene, alkylnaphthalenes, phenanthrene, and alkylphenanthrenes^17^. Molecular analysis of residual polycyclic aromatic hydrocarbons revealed that *M. sativa*+*P. tropica* treatment led to an almost complete degradation of these contaminants (Figure 3). These results are of great relevance considering the toxicity of these compounds. Among the United States Environmental Protection Agency’s 16 priority polycyclic aromatic hydrocarbons are naphthalene and phenanthrene^28^. Naphthalene is also classified as a group 2b carcinogen. In 2019, the European Chemical Agency added phenanthrene to the candidate list of substances of very high concern due to its very persistent, very bioaccumulative (vPvB) nature^29, 30^.

Finally, the relative abundances of *Paraburkholderia* in the residual uninoculated and inoculated soils were <1% and 5% respectively. This indicated that the inoculated bacteria thrived in the soils and were evidently responsible for the observed growth promotion of *M. sativa* and associated diesel fuel degradation. While many plant growth-promoting rhizobacteria have been experimented as inoculants for agricultural purposes, a major setback has always been the failure of inoculated microorganisms to effectively thrive against indigenous microbes^31^. Therefore, the results of this study will prove beneficial not only for environmental remediation but also for biotechnological applications in agriculture.

## MATERIALS AND METHODS

### Soil Sampling

A topsoil sample (10 g) was taken from a heavily polluted site in Wietze (52°39’0’’N, 09°50’0”E), Germany in November 2019 and transported to the laboratory on ice. Wietze is a site of historical petroleum production beginning in 1859^32^. Between 1900 and 1920, about 80% of German crude oil was produced in Wietze. The former oil field still contains petroleum seepages (Supplementary Figure S5) amidst a forested environment.

### Enrichment Cultures, Isolation of Single Bacterial Isolates and Growth Conditions

Approximately 1 g of the crude-oil-polluted soil was added to an Erlenmeyer flask (300 mL) containing 100 mL of liquid mineral medium (MM) composed of KH_2_PO_4_ (0.5 g/L), NaCl (0.5 g/L), NH_4_Cl (0.5 g/L). Sterile-filtered trace elements (1 mL/L)^33^, vitamin solution (1 mL/L)^33^ and MgSO_4_.7H_2_O (5 mL of a 100 mg/mL solution) were added to the MM, post MM-autoclaving. The pH was adjusted to 7.0. One mL of sterile-filtered diesel fuel was added to the flask as the sole carbon and energy source. The culture was grown at 30°C with shaking at 110 rpm (INFORS HT shaker, model CH-4103, Infors AG, Bottmingen, Switzerland) and subcultured after every five days. After three successive subculturing, the cells were plated on agar plates, after which diesel fuel was added to the plates using an airbrush. The plates were incubated at 30°C. After 48 hours, single colonies were transferred into separate flasks containing 100 mL liquid MM and 1 mL diesel fuel, and grown for five days. During the five-day period, bacterial growth in terms of optical density (OD_600_) was monitored every 12 hours using the UV/Visible spectrophotometer (Ultrospec 3000, Model 80-2106-20, Pharmacia Biotech, Cambridge England). Based on the OD_600_ values, three representative isolates were selected for whole genome studies.

### DNA Extraction

Microbial cells from approximately 30 mL of the cultures containing single isolates were harvested by centrifugation at 4000 x g for 10 min. DNA from the cell pellets were extracted using a MasterPure™ DNA Extraction kit (Epicentre^®^, Madison, USA) according to the manufacturer’s protocol. The DNA was used for Illumina-based whole genome sequencing. In addition, microbial cells from 30 mL of the cultures were harvested for Nanopore-based whole genome sequencing. Similarly, at the end of the experimental period, DNA from 100 mg of the inoculated and uninoculated residual soil samples were extracted using the PowerSoil^®^ DNA Extraction kit (Qiagen, Hilden, Germany).

### High-Throughput Sequencing

#### Genome sequencing and assembly

Genomes were sequenced at the Göttingen Genomics Laboratory, Germany. Short-reads were generated using a MiSeq sequencer and v3 chemistry (Illumina, San Diego, CA, USA), while long-reads were sequenced using a MinIon (Oxford Nanopore Technologies; Oxford, England). Reads were quality filtered with fastp version 0.20.1^34^ The leading 15cp were truncated from forward and reverse reads. Read shorter than 30bp were removed. Adaptor sequences were trimmed. Nanopore data were processed with porechop version 0.2.4^35^. Default parameters were used for all software unless otherwise specified. The quality of the processing was confirmed using FastQC version 0.91. Reads were assembled using Unicycler version 0.4.8^36^. Contigs shorter than 500bp were removed. Coverage information for each scaffold was determined using Bowtie 2 version 2.4.2^37^ and SAMtools version 1.11^38^. Genomes were classified taxonomically using GTDB-Tk version 1.0.2 and the Genome Taxonomy Database (release 86)^39, 40^.

#### Sequencing of bacterial 16S rRNA genes from residual soils

Bacterial 16S rRNA genes were amplified using the forward primer S-D-Bact-0341-b-S-17 (5’-CCT ACG GGN GGC WGC AG-3’)^41^ and the reverse primer S-D-Bact-0785-a-A-21 (5’-GAC TAC HVG GGT ATC TAA TCC-3’)^41^ containing Illumina Nextera adapters for sequencing. The PCR reaction (25 μL) contained 5 μL of five-fold Phusion HF buffer, 200 μM of each of the four deoxynucleoside triphosphates, 4 μM of each primer, 1 U of Phusion high fidelity DNA polymerase (Thermo Scientific, Waltham, MA, USA), and approximately 50 ng of the extracted DNA as a template. The negative controls were performed by using the reaction mixture without a template. The following thermal cycling scheme was used: initial denaturation at 98 °C for 30 s, 30 cycles of denaturation at 98 °C for 15 s, annealing at 53 °C for 30 s, followed by extension at 72 °C for 30 s. The final extension was carried out at 72 °C for 2 min. Obtained PCR products per sample were controlled for appropriate size and purified using the MagSi-NGS Plus kit according to the manufacturer’s protocol (Steinbrenner Laborsysteme GmbH, Germany). The quantification of the PCR products was performed using the Quant-iT dsDNA HS assay kit and a Qubit fluorometer, as recommended by the manufacturer (Thermo Scientific). The DNA samples were barcoded using the Nextera XT-Index kit (Illumina, San Diego, USA) and the Kapa HIFI Hot Start polymerase (Kapa Biosystems, USA). Sequencing was performed at the Göttingen Genomics Laboratory on an Illumina MiSeq Sequencing platform (paired end 2 × 300 bp) using the MiSeq Reagent kit v3, as recommended by the manufacturer (Illumina). All bacterial samples were sequenced on the same MiSeq run.

#### Processing of the 16S rRNA gene data

Trimmomatic version 0.39^42^ was initially used to truncate low quality reads if quality dropped below 12 in a sliding window of 4 bp. Datasets were subsequently processed with Usearch version 11.0.667^43^ as described in Wemheuer, Berkelmann, Wemheuer, Daniel, Vidal and Bisseleua Daghela^44^. In brief, paired end reads were merged and quality-filtered. Filtering included the removal of low-quality reads (maximum number of expected errors >2 and more than 1 ambitious base, respectively) and those shorter than 200 bp. Processed sequences of all samples were joined, dereplicated and clustered in zero-radius operational taxonomic units (zOTUs) using the UNOISE algorithm implemented in Usearch. A de novo chimera removal was included in the clustering step. Afterwards, zOTU sequences were taxonomically classified using the SINTAX algorithm against the SILVA database (SILVA SSURef 138 NR99). All non-bacterial zOTUs were removed based on their taxonomic classification. Subsequently, processed sequences were mapped on final zOTU sequences to calculate the distribution and abundance of each OTU in every sample. Richness and coverage based on the Chao1 richness estimator were estimated in R using the vegan package.

### Functional Analysis of the Genomes

#### Identification of CDSs involved in plant growth promoting and hydrocarbon-degrading processes

Coding DNA sequences of putative enzymes involved in both plant growth-promoting activities and hydrocarbon degradation were identified in the bacterial genomes by means of annotations with prodigal version 2.6.3^45^. Functional annotation was performed with diamond version v0.9.29^46^ and the KEGG database (October release 2018)^47^ The plant growthpromoting enzymes of interest include nif-specific regulatory protein, nitrogen fixation protein, electron transfer flavoprotein, acyl carrier protein, trehalose 6-phosphate phosphatase, phosphoglycolate phosphatase, phospholipase C, pyrroloquinoline quinone biosynthesis proteins B, D and E, pyrroloquinoline quinone-synthase, enterobactin (siderophore) exporter, and indoleacetamide hydrolase. Additionally, the genes responsible for bacterial chemotaxis, motility and root colonization were examined. These include the *che, wsp, flg, flh, fli* and *tad* genes. Similarly, the hydrocarbon-degrading enzymes of interest include long-chain alkane monooxygenase, cyclopentanol dehydrogenase, cyclohexanone monooxygenase, benzoate/toluate 1,2-dioxygenase, benzaldehyde dehydrogenase, dihydroxycyclohexadiene carboxylate dehydrogenase, catechol 1,2-dioxygenase, catechol 2,3-dioxygenase, muconate cycloisomerase, muconolactone D-isomerase, 3-oxoadipate enol-lactonase. On the basis of the following factors, *P. tropica* was selected for greenhouse-based rhizoremediation study: (1) the differences in the plant growth-promoting potentials of the different species as revealed by functional genomics, and (2) the differences in species’ tolerance to and utilization of diesel fuel hydrocarbons as shown by their growth rates (OD_600_) in the diesel-containing mineral medium.

#### Plant Growth and Bacterial Inoculation

The soil used for this experiment was “Primaster turf’, which is a mixture of screened sand, soil, and composted organics blended with an NPK fertiliser. The soil textural class was determined as sand (88.6% sand, 6.1% silt and 5.3% clay) with 12.5% organic matter content by loss on ignition, total nitrogen content of 0.15%, and a pH of 7.1. The soil was initially homogenized by sieving using a 2-mm mesh sieve to remove large particles. Diesel fuel-contaminated soil was prepared following the detailed procedure described in Eze, Thiel, Hose, George and Daniel^48^. Preliminary gas chromatography-mass spectrometry (GC-MS) analysis of the diesel fuel revealed the presence of fatty acid methyl esters (FAMEs) evidently from biodiesel (Supplementary Figure S6). Therefore, the soil was allowed to age for 7 days, so as to enable the removal of the FAMEs through natural attenuation by indigenous organisms. The resulting total petroleum hydrocarbons in the soil after ageing (designated as time T0) was determined using GC-MS. Viable seeds of *Medicago sativa* L. were placed in pots (3 seeds per pot) containing 150 g of the aged diesel fuel-contaminated soils.

The contaminated soil was treated with the following: (1) *M. sativa*; (2) *P. tropica*; (3) *M. sativa*+*P. tropica*. An unplanted and uninoculated soil served as the control. Since the goal of the study was to assess the effectiveness of each treatment for hydrocarbon degradation, the soil used for the entire experiment was the diesel fuel-contaminated soil described above. The microbial consortium used was harvested from the culture by centrifugation at 4000 x g for 10 min, washed twice in mineral medium and concentrated to OD_600_ = 1.650. The same amount of cells were applied to both the “*P. tropica*” and the “*M. sativa*+*P. tropica*” treatments. For the *M. sativa*+*P. tropica* treatment, the cells (at OD_600_ = 1.650) were inoculated to the base of one-week-old *M. sativa* seedlings at the depth of 1.5 cm below-ground level. The whole experiment was performed in triplicate, and pots were watered with 90 mL sterile water every three days for the first two weeks. After that, the planted pots were watered with 90 mL sterile water every two days to compensate for the water needs of *M. sativa* plants. To assess the effect of bacterial inoculation on growth rate, shoot heights (in mean values of plants per pot) attained with time were taken every two weeks. Plant height was measured from the shoot tip to the base of stem^15, 49^ After 60 days, each plant was harvested, washed under tap water, oven-dried at 70°C until constant weights were achieved, and then their dry biomass weights were recorded.

### Geochemical Assessment of Microbial-Enhanced Bioremediation

#### Extraction of residual hydrocarbons

After 60 days, *M. sativa* plants were harvested from the pots. The soil samples from each pot were first manually homogenized. For hydrocarbon analyses, 1 g of the ground freeze-dried soils were further homogenized with a small amount of sodium sulfate (Na_2_SO_4_) and transferred into a Teflon microwave digestion vessel. The samples were solvent extracted twice with fresh 2.5 mL n-hexane in a microwave device (Mars Xpress, CEM; 1600W, 100°C, 20 min). For reference, 2.5 μL diesel fuel (density = 0.82 g/mL) were dissolved in 5 mL *n*-hexane instead of 1 g soil sample. The extracts were combined into 7 mL vials and topped to 5 mL with *n*-hexane. A 1 mL aliquot (20%) of each extract was pipetted into a 2 mL autosampler vial, and 20 μL *n*-icosane D42 [200 mg/L] was added as an internal quantification standard.

#### Molecular analysis of biodegradation

GC-MS analyses of the samples were performed using a Thermo Scientific Trace 1300 Series GC coupled to a Thermo Scientific Quantum XLS Ultra MS. The GC capillary column was a Phenomenex Zebron ZB–5MS (30 m, 0.1 μm film thickness, inner diameter 0.25 mm). Compounds were transferred splitless to the GC column at an injector temperature of 300°C. Helium was used as the carrier gas at a flow rate of 1.5 mL/min. The GC temperature program was as follows: 80°C (hold 1 min), 80°C to 310°C at 5°C/min (hold 20 min). Electron ionization mass spectra were recorded at 70eV electron energy in full scan mode (mass range m/z 50–600, scan time 0.42 s). Peak areas were integrated using Thermo Xcalibur software version 2.2 (Thermo Fisher Scientific Inc., USA).

#### Biodegradation parameters

To assess the nature and extent of biodegradation in the different treatments, the ratios of *n*-hexadecane (nC16), *n*-heptadecane (nC17) and *n*-octadecane (nC18) versus the more refractory isoprenoid hydrocarbons nor-pristane (2,6,10-trimethylpentadecane, nor-Pr), pristane (2,6,10,14-tetramethylpentadecane, Pr) and phytane (2,6,10,14-tetramethylhexadecane, Ph) were calculated. As an additional parameter, the relative abundance of total petroleum hydrocarbons (TPH) versus the unresolved complex mixture (UCM, often referred to as the “hump”, a typical indicator of biodegradation^50^) was determined.

#### Statistical Analyses

All statistical analysis were performed using R^51^. One-way analysis of variance (ANOVA) was used to compare the mean dry biomass of *M. sativa* and *M. sativa*+*P. tropica* treatments. The normality and homogeneity of variances were tested by the Shapiro-Wilk’s test^52^ and the Levene’s test^53^ respectively. Relative growth rates of plants under different treatments were determined following the method of Eze, George and Hose^15^. This method involved the assessment of growth rate in terms of mean shoot height per pot attained with time. This approach eliminates the biases associated with destructive harvesting methods^54^. The logistic model was used for the statistical analysis of relative growth rate^55, 56^. The lower asymptote *c*was fixed at 0 since height at time *t_0_* is 0, resulting in a 3-parameter logistic model.

Similarly, comparisons of soil hydrocarbon concentrations before (T0) and after (T60), and between treatments were made using one-way ANOVA, followed by Tukey’s all-pairwise comparisons. In all cases, the normality of variances was tested by the Shapiro-Wilk’s method^52^, and homogeneity of variances was tested using the Levene’s test^53^. The significance level (nominally 0.05) was adjusted for multiple comparisons using the Holm method^57, 58^.

### Author contributions

Conceptualization and design: MOE, GCH, SCG and RD. Planning and implementation: MOE and RD. Experiments and bioinformatics analyses: MOE. Geochemical analysis: MOE and VT. Writing – original draft: MOE. Writing – review and editing: VT, GCH, SCG and RD. Supervision: GCH, SCG and RD.

## Acknowledgements

The authors would like to thank the Commonwealth Government of Australia and the German Academic Exchange Service (DAAD) for supporting this research project by providing M.O.E. with an international Research Training Program (iRTP) Scholarship and DAAD Scholarship (Allocation Numbers: 2017561 and 91731339 respectively). This publication was supported financially by the Open Access Grant Program of the German Research Foundation (DFG) and the Open Access Publication Fund of the University of Goettingen. We also thank Dr. Anja Poehlein and Melanie Heinemann for the assistance during sequencing.

## References

1. Bragg, J.R., Prince, R.C., Harner, E.J. & Atlas, R.M. Effectiveness of bioremediation for the Exxon Valdez oil spill. Nature 368, 413–418 (1994).

2. Abbriano, R. et al. Deepwater Horizon oil spill: a review of the planktonic response. Oceanography 24, 294–301 (2011).

3. Hemmer, M.J., Barron, M.G. & Greene, R.M. Comparative toxicity of eight oil dispersants, Louisiana sweet crude oil (LSC), and chemically dispersed LSC to two aquatic test species. Environ. Toxicol. Chem. 30, 2244–2252 (2011).

4. Duffy, J.J., Peake, E. & Mohtadi, M.F. Oil spills on land as potential sources of groundwater contamination. Environ. Int. 3, 107–120 (1980).

5. Azubuike, C.C., Chikere, C.B. & Okpokwasili, G.C. Bioremediation techniques–classification based on site of application: principles, advantages, limitations and prospects. World J. Microbiol. Biotechnol. 32, 180 (2016).

6. USEPA, EPA/600/R-99/107. Introduction to phytoremediation. United States Environmental Protection Agency (2000).

7. USEPA, EPA 542-R-01-006. Brownfields technology primer: selecting and using phytoremediation for site cleanup. United States Environmental Protection Agency (2001).

8. Meagher, R.B. Phytoremediation of toxic elemental and organic pollutants. Current Opin. Plant Biol. 3, 153–162 (2000).

9. Bevivino, A., Dalmastri, C., Tabacchioni, S. & Chiarini, L. Efficacy of *Burkholderia cepacia* MCI 7 in disease suppression and growth promotion of maize. Biology and Fertility of Soils 31, 225–231 (2000).

10. Rohrbacher, F. & St-Arnaud, M. Root exudation: the ecological driver of hydrocarbon rhizoremediation. Agronomy 6, 19 (2016).

11. Garrido-Sanz, D. et al. Metagenomic insights into the bacterial functions of a diesel-degrading consortium for the rhizoremediation of diesel-polluted soil. Genes 10, 456 (2019).

12. Glick, B.R. Phytoremediation: synergistic use of plants and bacteria to clean up the environment. Biotechnol. Adv. 21, 383–393 (2003).

13. Dias, G.M. et al. Comparative genomics of *Paraburkholderia kururiensis* and its potential in bioremediation, biofertilization, and biocontrol of plant pathogens. MicrobiologyOpen 8, e00801 (2019).

14. García, S.S. et al. *Paraburkholderia tropica* as a plant-growth–promoting bacterium in barley: characterization of tissues colonization by culture-dependent and - independent techniques for use as an agronomic bioinput. Plant Soil (2019).

15. Eze, M.O., George, S.C. & Hose, G.C. Dose-response analysis of diesel fuel phytotoxicity on selected plant species. Chemosphere 263, 128382 (2021).

16. Eze, M.O. & George, S.C. Ethanol-blended petroleum fuels: implications of co-solvency for phytotechnologies. RSC Adv. 10, 6473–6481 (2020).

17. ATSDR. Toxicological profile for fuel oil. Agency for Toxic Substances and Disease Registry, United States (1995).

18. Esmaeel, Q. et al. *Paraburkholderia phytofirmans* PsJN-plants interaction: from perception to the induced mechanisms. Front. Microbiol. 9, 2093 (2018).

19. Böhm, M., Hurek, T. & Reinhold-Hurek, B. Twitching motility is essential for endophytic rice colonization by the N2-fixing endophyte *Azoarcus* sp. strain BH72. Mol. Plant Microbe Interact. 20, 526–533 (2007).

20. Brencic, A. & Winans, S.C. Detection of and response to signals involved in host-microbe interactions by plant-associated bacteria. Microbiol. Mol. Biol. Rev. 69, 155–194 (2005).

21. Chen, Y. et al. Comparative genomic analysis and phenazine production of Pseudomonas chlororaphis, a plant growth-promoting rhizobacterium. Genom. Data 4, 33–42 (2015).

22. Haq, I.U., Graupner, K., Nazir, R. & van Elsas, J.D. The genome of the fungal-interactive soil bacterium *Burkholderia terrae* BS001—a plethora of outstanding interactive capabilities unveiled. Genome Biol. Evol. 6, 1652–1668 (2014).

23. Shen, X., Hu, H., Peng, H., Wang, W. & Zhang, X. Comparative genomic analysis of four representative plant growth-promoting rhizobacteria in *Pseudomonas*. BMC Genom. 14, 271 (2013).

24. Lee, Y., Lee, Y. & Jeon, C.O. Biodegradation of naphthalene, BTEX, and aliphatic hydrocarbons by *Paraburkholderia aromaticivorans* BN5 isolated from petroleum-contaminated soil. Sci. Rep. 9, 860 (2019).

25. Gemmell, R.T. & Knowles, C.J. Utilisation of aliphatic compounds by acidophilic heterotrophic bacteria. The potential for bioremediation of acidic wastewaters contaminated with toxic organic compounds and heavy metals. FEMS Microbiol. Lett. 192, 185–190 (2000).

26. Röling, W.F.M., Ortega-Lucach, S., Larter, S.R. & Head, I.M. Acidophilic microbial communities associated with a natural, biodegraded hydrocarbon seepage. J. Appl. Microbiol. 101, 290–299 (2006).

27. Giovanella, P. et al. Metal and organic pollutants bioremediation by extremophile microorganisms. J. Hazard. Mater. 382, 121024 (2020).

28. Bojes, H.K. & Pope, P.G. Characterization of EPA’s 16 priority pollutant polycyclic aromatic hydrocarbons (PAHs) in tank bottom solids and associated contaminated soils at oil exploration and production sites in Texas. Regul. Toxicol. Pharmacol. 47, 288–295 (2007).

29. ECHA, ECHA/PR/19/01 The candidate list of substances of very high concern (SVHCs). European Chemical Agency (2019).

30. Loibner, A.P., Szolar, O.H.J., Braun, R. & Hirmann, D. Toxicity testing of 16 priority polycyclic aromatic hydrocarbons using Lumistox^®^. Environ. Toxicol. Chem. 23, 557–564 (2004).

31. Bashan, Y., de-Bashan, L.E., Prabhu, S.R. & Hernandez, J.-P. Advances in plant growth-promoting bacterial inoculant technology: formulations and practical perspectives (1998–2013). Plant Soil 378, 1–33 (2014).

32. Craig, J., Gerali, F., MacAulay, F. & Sorkhabi, R. The history of the European oil and gas industry (1600s-2000s). Geol. Soc. Spec. Publ. 465, 1 (2018).

33. Atlas, R.M. Handbook of microbiological media, 4th edition. CRC Press, Boca Raton, Florida (2010).

34. Chen, S., Zhou, Y., Chen, Y. & Gu, J. fastp: an ultra-fast all-in-one FASTQ preprocessor. Bioinformatics 34, i884–i890 (2018).

35. Wick, R. Porechop, Github (https://github.com/rrwick/Porechop) (2017).

36. Wick, R.R., Judd, L.M., Gorrie, C.L. & Holt, K.E. Unicycler: Resolving bacterial genome assemblies from short and long sequencing reads. PLOS Comput. Biol. 13, e1005595 (2017).

37. Langmead, B. & Salzberg, S.L. Fast gapped-read alignment with Bowtie 2. Nat. Methods 9, 357–359 (2012).

38. Li, H. et al. The sequence alignment/map format and SAMtools. Bioinformatics 25, 2078–2079 (2009).

39. Chaumeil, P.-A., Mussig, A.J., Hugenholtz, P. & Parks, D.H. GTDB-Tk: a toolkit to classify genomes with the Genome Taxonomy Database. Bioinformatics 36, 1925–1927 (2019).

40. Parks, D.H. et al. Selection of representative genomes for 24,706 bacterial and archaeal species clusters provide a complete genome-based taxonomy. bioRxiv, 771964 (2019).

41. Klindworth, A. et al. Evaluation of general 16S ribosomal RNA gene PCR primers for classical and next-generation sequencing-based diversity studies. Nucleic Acids Res. 41, e1–e1 (2013).

42. Bolger, A.M., Lohse, M. & Usadel, B. Trimmomatic: a flexible trimmer for Illumina sequence data. Bioinformatics 30, 2114–2120 (2014).

43. Edgar, R.C. Search and clustering orders of magnitude faster than BLAST. Bioinformatics 26, 2460–2461 (2010).

44. Wemheuer, F. et al. Agroforestry management systems drive the composition, diversity, and function of fungal and bacterial endophyte communities in *Theobroma cacao* leaves. Microorganisms 8 (2020).

45. Hyatt, D. et al. Prodigal: prokaryotic gene recognition and translation initiation site identification. BMC Bioinform. 11, 119–119 (2010).

46. Buchfink, B., Xie, C. & Huson, D.H. Fast and sensitive protein alignment using DIAMOND. Nat. Methods 12, 59–60 (2015).

47. Kanehisa, M. & Goto, S. KEGG: kyoto encyclopedia of genes and genomes. Nucleic Acids Res. 28, 27–30 (2000).

48. Eze, M.O., Thiel, V., Hose, G.C., George, S.C. & Daniel, R. Metagenomic insight into the plant growth-promoting potential of a diesel-degrading bacterial consortium for enhanced rhizoremediation application. bioRxiv, 2021.2003.2026.437261 (2021).

49. Chen, S., Li, J., Fritz, E., Wang, S. & Hüttermann, A. Sodium and chloride distribution in roots and transport in three poplar genotypes under increasing NaCl stress. For. Ecol. Manag. 168, 217–230 (2002).

50. Peters, K.E., Walters, C.C. & Moldowan, J.M. The Biomarker Guide: Volume 2: Biomarkers and Isotopes in Petroleum Systems and Earth History. Cambridge University Press (2004).

51. R Core Team. R: a languange and environment for statistical computing. R Foundation for Statistical Computing, Vienna, Austria (2018).

52. Ghasemi, A. & Zahediasl, S. Normality tests for statistical analysis: a guide for non-statisticians. Int. J. Endocrinol. Metab. 10, 486–489 (2012).

53. Levene, H. Robust tests for equality of variances, in: Contributions to Probability and Statistics; Essays in Honor of Harold Hotelling (eds. O. Ingram & H. Harold) 278–292. Stanford University Press, Stanford, California (1960).

54. Hoffmann, W.A. & Poorter, H. Avoiding bias in calculations of relative growth rate. Ann. Bot. 90, 37–42 (2002).

55. Gregorczyk, A. A logistic function—its application to the description and prognosis of plant growth. Acta Soc. Bot. Pol. 60, 67–76 (1991).

56. Szparaga, A. & Kocira, S. Generalized logistic functions in modelling emergence of *Brassica napus* L. PLOS ONE 13, e0201980 (2018).

57. Lee, S. & Lee, D.K. What is the proper way to apply the multiple comparison test? Korean J. Anesthesiol. 71, 353–360 (2018).

58. Holm, S. A simple sequentially rejective multiple test procedure. Scand. J. Stat. 6, 65–70 (1979).

