## Supplementary Materials for "Exploiting synergistic interactions of *Medicago sativa* L. and *Paraburkholderia tropica* for enhanced biodegradation of diesel fuel hydrocarbons"

**Supplementary Table S1**. Sequencing results for the three genomes

| Genomes | Size (bp) | Raw reads | Fastp | | Scaffolds | Scaffolds (>500 bp) | Coverage |
|---|---|---|---|---|---|---|---|
| | | | Paired reads | Unpaired reads | | | |
| *Acidocella facilis* | 4,079,951 | 1,234,060 | 1,119,120 | 52,554 | 216 | 181 | 44.4 |
| *Burkholderia* sp. | 7,741,599 | 813,998 | 636,137 | 79,504 | 263 | 235 | 13.1 |
| *Paraburkholderia tropica* | 8,454,837 | 2,180,716 | 1,695,018 | 223,965 | 96 | 81 | 31.8 |
| **Total** | | **4,228,774** | **3,450,275** | **356,023** | **575** | **497** | **Avg: 30x** |

**Supplementary Table S2**. Plant growth promoting genes and number of CDSs for the three genomes

| Gene | Genomes | | | Enzyme name |
|---|---|---|---|---|
| | **1** | **2** | **3** | |
| ***Plant growth promotion*** | | | | |
| **Nitrogen metabolism** | | | | |
| *nifA* | 0 | 5 | 2 | Nif-specific regulatory protein |
| *nifU* | 1 | 1 | 1 | Nitrogen fixation protein NifU and related proteins |
| *nifQ* | 0 | 1 | 1 | Nitrogen fixation protein NifQ |
| *fixA* | 2 | 5 | 5 | Electron transfer flavoprotein beta subunit |
| *fixB* | 2 | 4 | 5 | Electron transfer flavoprotein alpha subunit |
| *fixL* | 3 | 1 | 11 | Two-component system, sensor kinase FixL |
| *fixJ* | 3 | 2 | 11 | Two-component system, response regulator FixJ |
| **Phosphate solubilization** | | | | |
| *acpS* | 1 | 1 | 1 | Holo-[acyl-carrier protein] synthase [EC:2.7.8.7] |
| *acpP* | 1 | 2 | 4 | Acyl carrier protein |
| *serB* | 1 | 1 | 1 | Phosphoserine phosphatase [EC:3.1.3.3] |
| *otsB* | 4 | 2 | 3 | Trehalose 6-phosphate phosphatase [EC:3.1.3.12] |
| *gph* | 3 | 6 | 6 | Phosphoglycolate phosphatase [EC:3.1.3.18] |
| *phoD* | 0 | 1 | 1 | Alkaline phosphatase D [EC:3.1.3.1] |
| *plc* | 1 | 10 | 9 | Phospholipase C [EC:3.1.4.3] |
| *glpR* | 4 | 4 | 6 | Glycerol-3-phosphate regulon repressor |
| **Pyrroloquinoline quinone synthesis** | | | | |
| *pqqB* | 1 | 1 | 1 | Pyrroloquinoline quinone biosynthesis protein B |
| *pqqC* | 1 | 1 | 1 | Pyrroloquinoline-quinone synthase [EC:1.3.3.11] |
| *pqqD* | 2 | 1 | 1 | Pyrroloquinoline quinone biosynthesis protein D |
| *pqqE* | 1 | 1 | 1 | Pyrroloquinoline quinone biosynthesis protein E |
| **Siderophore transport** | | | | |
| *entS* | 0 | 1 | 2 | Enterobactin (siderophore) exporter |
| **Indoleacetic acid synthesis** | | | | |
| *iaaH* | 0 | 0 | 2 | Indoleacetamide hydrolase [EC:3.5.1.-] |
| **Total** | **31** | **51** | **75** | |
| ***Bacterial chemotaxis, motility, root nodulation and colonization*** | | | | |
| *cheABCDVWXYZ* | 12 | 13 | 20 | Two-component chemotaxis protein |
| *wspBDEF* | 0 | 4 | 4 | Two-component chemotaxis protein |
| *mcp, tsr, tar, trg, tap, wspA* | 7 | 11 | 47 | Methyl-accepting chemotaxis protein |
| *flg, flh, fli* | 43 | 48 | 52 | Flagellar proteins |
| *nodD* | 1 | 3 | 2 | Nod-box dependent transcriptional activator |
| *tadB* | 1 | 2 | 2 | Tight adherence protein B |
| *tadC* | 1 | 2 | 2 | Tight adherence protein C |
| **Total** | **65** | **83** | **129** | |

Genome 1: *Acidocella facilis;* Genome 2: *Burkholderia* sp.; Genome 3: *Paraburkholderia tropica*.

**Supplementary Table S3**. Selected genes putatively involved in hydrocarbon degradation and number of CDSs

| Gene | Genomes | | | Enzyme name |
|---|---|---|---|---|
| | **1** | **2** | **3** | |
| ***Aliphatic hydrocarbon degradation*** | | | | |
| **n-Alkanes** | | | | |
| *alkM* | 2 | 1 | 0 | Alkane 1-monooxygenase |
| *alkR* | 1 | 2 | 2 | Alkane utilization regulator |
| *ladA* | 3 | 6 | 6 | Long-chain alkane monooxygenase |
| *prmB* | 1 | 0 | 1 | Propane monooxygenase reductase component |
| **Cycloalkanes** | | | | |
| *cpnA* | 0 | 4 | 2 | Cyclopentanol dehydrogenase |
| *chnB* | 0 | 2 | 2 | Cyclohexanone monooxygenase |
| *gnl* | 3 | 1 | 5 | Gluconolactonase |
| *chnD* | 0 | 0 | 1 | 6-Hydroxyhexanoate dehydrogenase |
| ***Aromatic hydrocarbon degradation*** | | | | |
| *tmoF* | 0 | 0 | 2 | Toluene monooxygenase |
| *pobA* | 1 | 1 | 1 | p-Hydroxybenzoate 3-monooxygenase |
| *adhPE* | 7 | 12 | 13 | Alcohol dehydrogenase |
| *benABC* | 0 | 6 | 6 | Benzoate/toluate 1,2-dioxygenase |
| *benD* | 1 | 1 | 1 | Dihydroxycyclohexadiene carboxylate dehydrogenase |
| *pcaGH* | 4 | 2 | 2 | Protocatechuate 3,4-dioxygenase |
| *etbD* | 0 | 1 | 1 | 2-Hydroxy-6-oxo-octa-2,4-dienoate hydrolase |
| *catA* | 0 | 3 | 2 | Catechol 1,2-dioxygenase |
| *catB* | 0 | 2 | 2 | Muconate cycloisomerase |
| *catC* | 0 | 2 | 2 | Muconolactone D-isomerase |
| *pcaDL* | 6 | 8 | 8 | 3-Oxoadipate enol-lactonase |
| *dmpB* | 2 | 2 | 1 | Catechol 2,3-dioxygenase |
| *praC* | 1 | 4 | 4 | 4-Oxalocrotonate tautomerase |
| *mhpD* | 0 | 2 | 2 | 2-Keto-4-pentenoate hydratase |
| *mhpE* | 0 | 1 | 1 | 4-Hydroxy 2-oxovalerate aldolase |

Genome 1: *Acidocella facilis*; Genome 2: *Burkholderia* sp.; Genome 3: *Paraburkholderia tropica*.

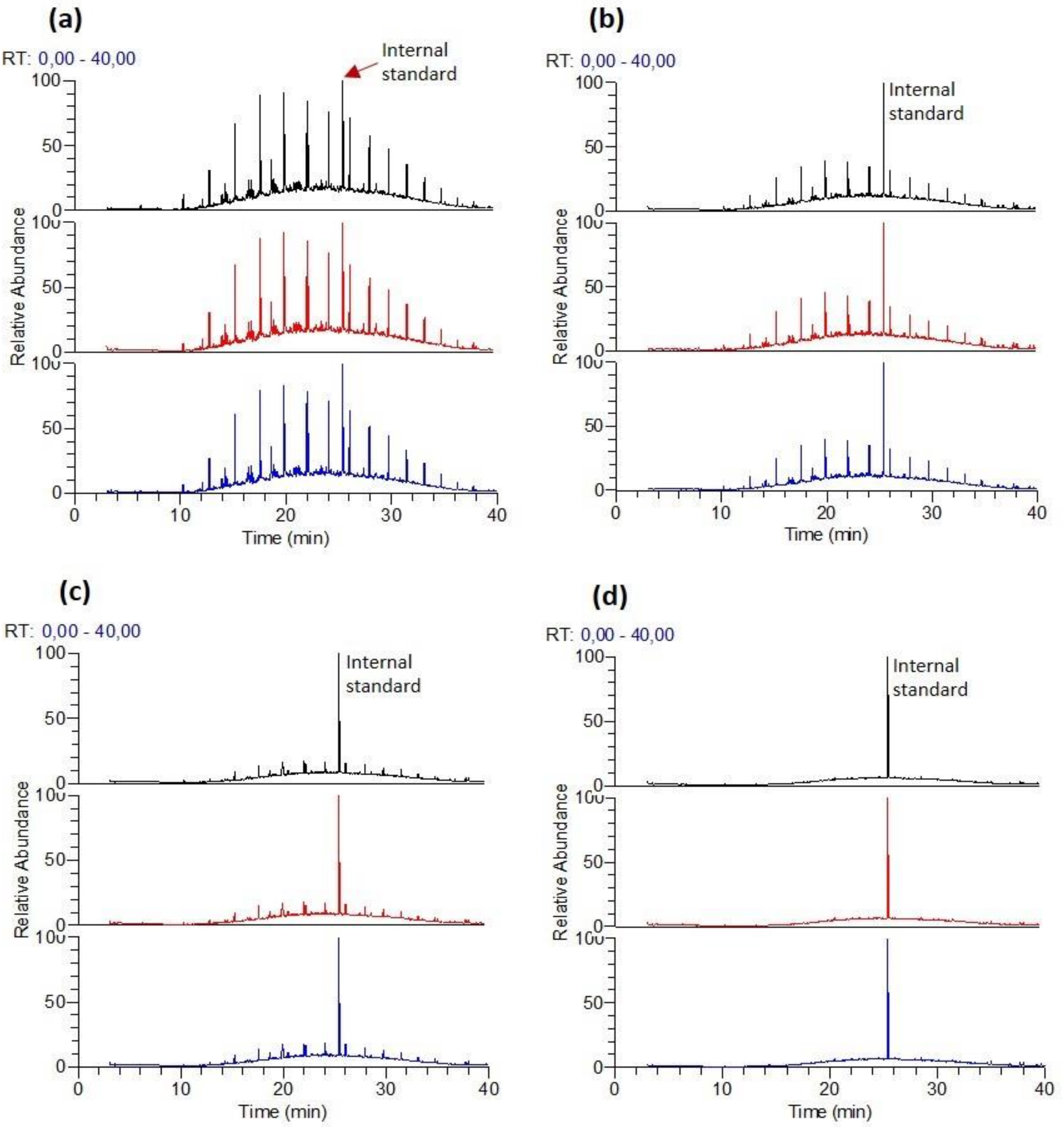


**Supplementary Figure S1**. Partial total ion chromatograms showing replicates of (a) Soil at Tf, (b) Soil+*M.sativa*, (c) Soil+*P.tropica*, and (d) Soil+*M.sativa*+*P.tropica*. Refer to Figure 2 in the main text for sample descriptions.

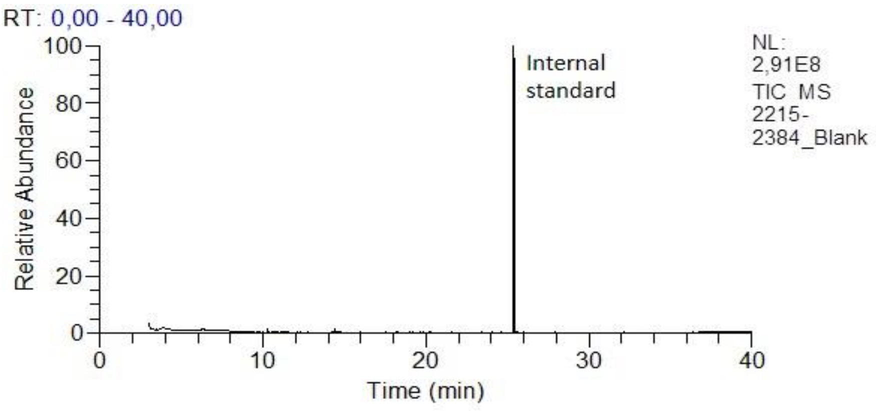


**Supplementary Figure S2**. Partial total ion chromatogram of preparation blank used to control cross contamination.

**Supplementary Table S4**. Statistical parameters for Tukey's test showing significant differences in the mean values of residual hydrocarbons between treatments.

| Linear hypothesis (Tukey contrasts) | Estimate | Std. Error | *t* value | Pr(>\|t\|) |
|---|---|---|---|---|
| Soil at T60 – Soil at T0 == 0 | -2.0300 | 0.0738 | -27.50 | 9.37e-10 *** |
| Soil+*M.sativa* – Soil at T0 == 0 | -2.9700 | 0.0738 | -40.23 | 2.15e-11 *** |
| Soil+*P.tropica* – Soil at T0 == 0 | -3.5467 | 0.0738 | -48.05 | 3.68e-12 *** |
| Soil+*M.sativa*+*P.tropica* – Soil at T0 == 0 | -3.9833 | 0.0738 | -53.96 | 1.16e-12 *** |
| Soil+*M.sativa* – Soil at T60 == 0 | -0.9400 | 0.0738 | -12.73 | 1.67e-06 *** |
| Soil+*P.tropica* – Soil at T60 == 0 | -1.5167 | 0.0738 | -20.55 | 1.65e-08 *** |
| Soil+*M.sativa*+*P.tropica* – Soil at T60 == 0 | -1.9533 | 0.0738 | -26.46 | 1.37e-09 *** |
| Soil+*P.tropica* – Soil+*M.sativa* == 0 | -0.5767 | 0.0738 | -7.81 | 1.45e-04 *** |
| Soil+*M.sativa*+*P.tropica* – Soil+*M.sativa* == 0 | -1.0133 | 0.0738 | -13.73 | 8.17e-07 *** |
| Soil+*M.sativa*+*P.tropica* – Soil+*P.tropica* == 0 | -0.4367 | 0.0738 | -5.92 | 0.0015 ** |

Significant codes: 0 '***' 0.001 '**' 0.01 '*' 0.05 '.' 0.1 ' ' 1. (Adjusted *p* values reported -- Holm method).

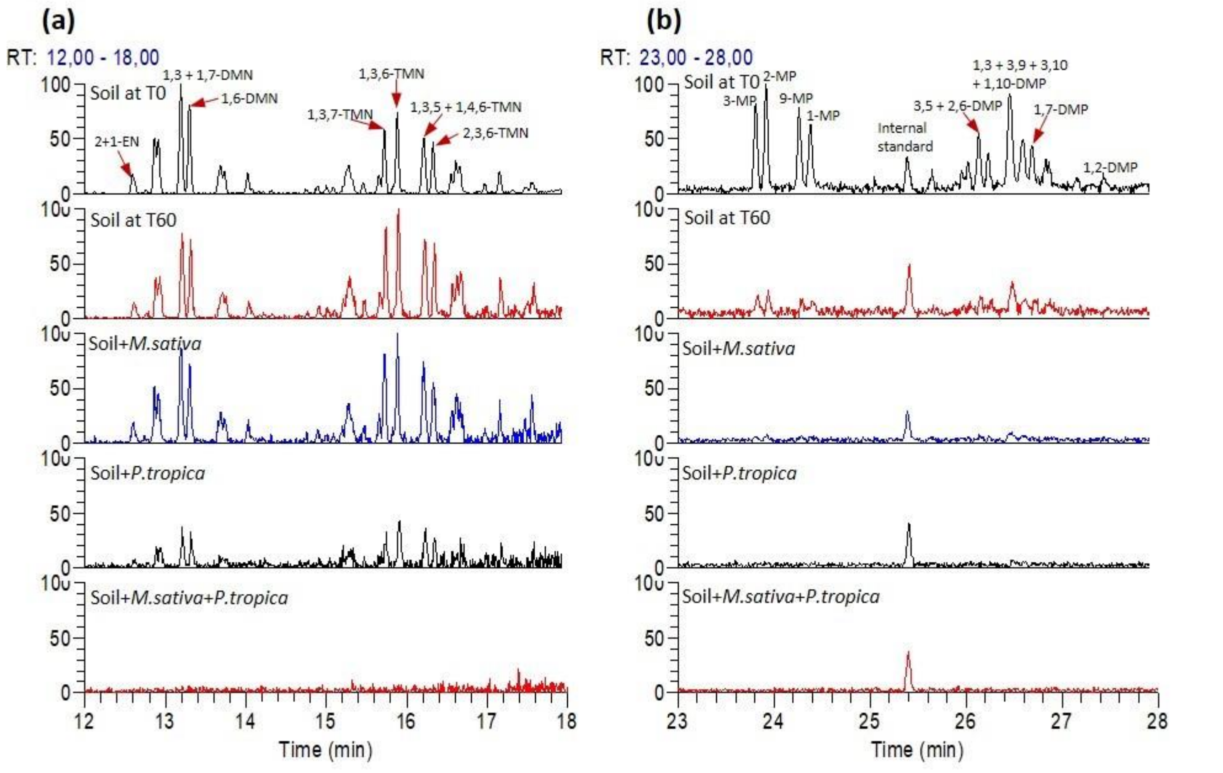


**Supplementary Figure S3**. Partial (a) *m/z* 156 + 170 and (b) *m/z* 192 + 206 GC-MS chromatograms of the different treatments showing differential biodegradation of naphthalenes and phenanthrenes. EN: ethylnaphthalene; DMN: dimethylnaphthalene; MP: methylphenanthrene; DMP: dimethylphenanthrene. Numbers denote positions of alkylation.

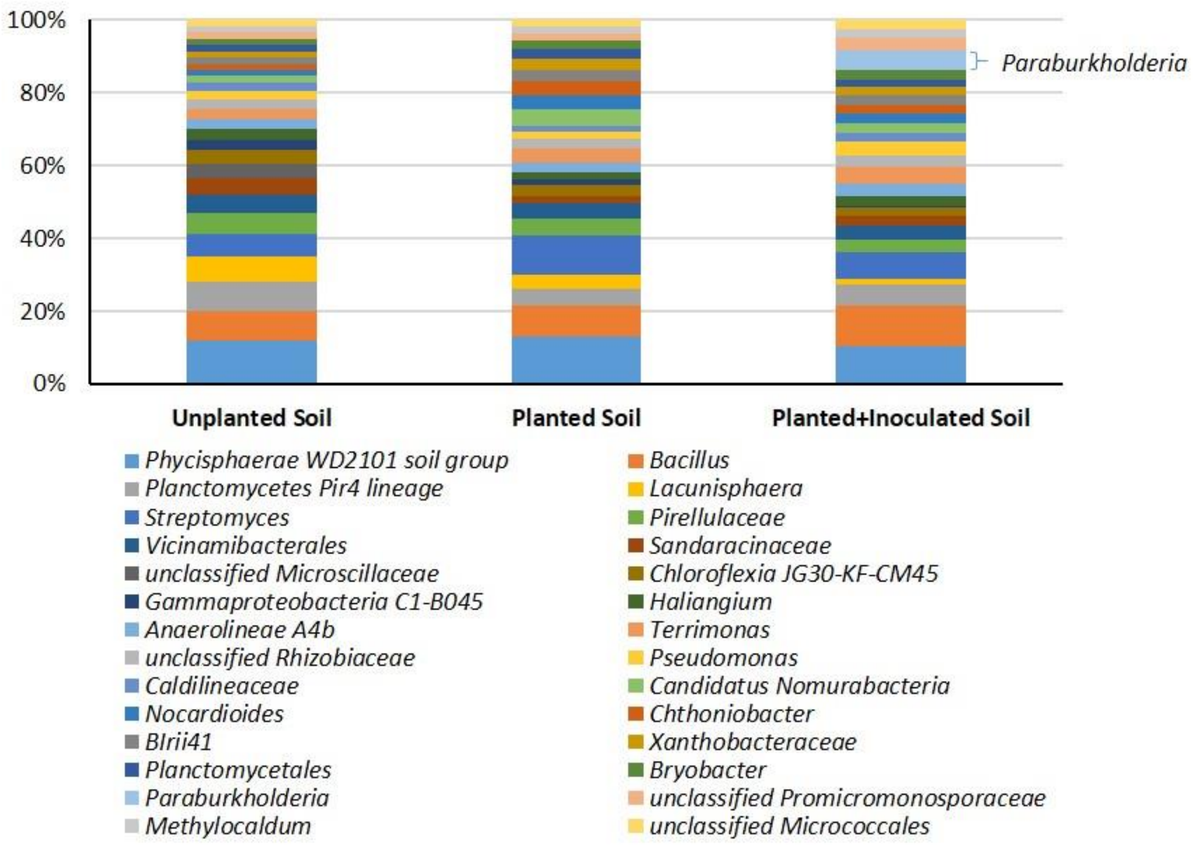


**Supplementary Figure S4**. Relative abundances of bacterial populations in the residual soils based on 16S rRNA gene amplicon data. Only taxa with relative abundance of ≥ 1% are presented.

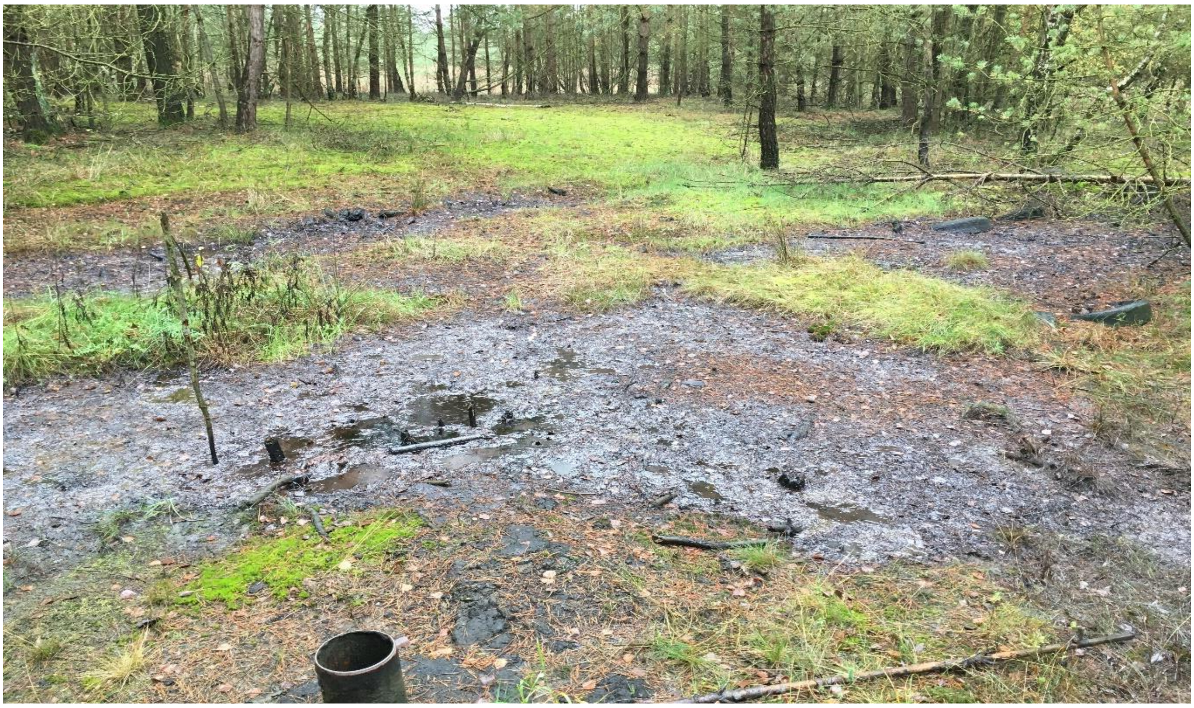

**Supplementary Figure S5.** Picture of the sampling site in Wietze, Germany.

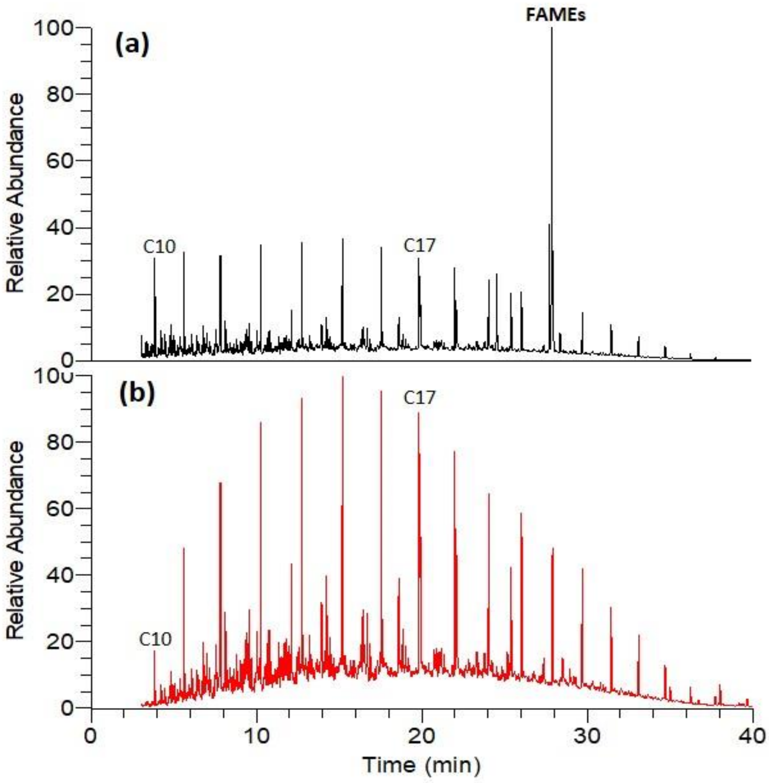


**Supplementary Figure S6**. Partial total ion chromatograms of (a) pure diesel fuel prior to soil spiking and ageing showing the presence of fatty acid methyl esters (FAMEs) from a biodiesel component, and (b) extracted diesel fuel after ageing showing the absence of FAMEs.
